# F_1_F_0_-ATP-synthase subunit *b* head-to-head interactions shape intracellular membranes in *Escherichia coli*

**DOI:** 10.64898/2026.03.03.708934

**Authors:** Mukul S. Kareya, Jorge Royes, Federica Angius, Florence Szczepaniak, Francesco Boldrin, Marion Hamon, Pauline Talbot, Oana Ilioaia, Céline Madigou, Christophe Tribet, François Dehez, Florent Waltz, Francesca Zito, Bruno Miroux

**Affiliations:** Université Paris Cité, CNRS, Biochimie des Protéines Membranaires, F-75005 Paris, France; CNRS, LPCT, Université de Lorraine, F-54000 Nancy, France; Department of Biology, University of Padova, Padova, Italy; Département de Chimie, École Normale Supérieure, PSL University, CNRS, Sorbonne Université, Paris, France; Biozentrum, University of Basel, Basel, Switzerland; Department of Chemistry, Biochemistry, Johannes Gutenberg University Mainz, Germany; iSpheres, Montpellier, Occitanie, France; Excelya, Boulogne Billancourt, France; Department of Biological Sciences, Lehigh University, United States; EverZom, Paris, France; Biologie Computationnelle et Quantitative, UMR 7238 CNRS - Sorbonne Université, Paris, France; ESPCI Paris, Paris, France

## Abstract

Intracellular membrane (ICM) compartments in bacteria remain poorly understood. In *Escherichia coli*, ICMs form upon the overproduction of certain membrane proteins. Here, we investigate at the molecular level how overproduction of subunit *b* of the F_1_F_o_-ATP synthase complex shapes ICMs. Through a combination of subunit *b* deletions, electron microscopy, and biochemical analyses, we demonstrate that ICMs network structuration is mediated by the C-terminal domain of subunit *b*. High-resolution cryo-electron tomography and molecular dynamics simulations reveal that this domain forms tetrameric assemblies, bridging membranes and maintaining a regular 26 nm spacing. Transcriptomic and proteomic analyses further show that the cell responds to ICMs formation by upregulating energy metabolism genes and the ESCRT-III homolog PspA, while downregulating ribosomal and translational machinery.

These results uncover a novel mechanism by which the C-terminal domain of subunit *b* drives the structuring of intracellular membranes. Additionally, they highlight how *E. coli* adapts to membrane stress by overexpressing membrane-remodeling proteins, a response that shares similarities with intracellular membrane homeostasis mechanisms in photosynthetic bacteria and unicellular algae.

## 2 Introduction

A wide variety of bacteria exhibit complex, spatially organized subcellular compartments such as intracytoplasmic membranes (ICMs)[1, 2]. Bacterial homologs of the mitochondrial crista-developing protein Mic60[2], the dynamin-like protein[3], and the eukaryotic and archaeal endosomal sorting complexes required for transport (ESCRT-III) superfamily[1] play an important role in membrane remodeling and biogenesis. The bacterial proteins PspA and Vipp1/IM30 share this remodeling function in bacteria[4] and cyanobacteria[5], respectively, as well as in the thylakoid membranes of chloroplasts. They form oligomeric assemblies that interact with membrane phospholipids to bend and increase membrane curvature during fusion, fission, and repair processes[4].

Most gram-negative species, such as *Escherichia coli*, lack intracytoplasmic membranes; however, ICMs formation in *E. coli* has been observed upon overexpression of a few membrane proteins[6]. To date, the underlying molecular mechanisms driving ICMs formation remain elusive[7, 8]. One example of ICMs formation is that mediated by the F_1_F_0_-ATP synthase protein complex (Fig. 1, Panel a), which synthesizes ATP from ADP using the proton-motive force. The F_0_ domain is composed of the a, *b*, and c subunits and rotates upon proton translocation through the inner membrane. The *γ* subunit connects the F_0_ domain to the catalytic F_1_ domain. Rotation of the *γ* subunit induces sequential conformational changes in the catalytic sites of the F_1_ *β*-subunit, allowing ADP binding, ATP synthesis, and ATP release.

**Fig. 1.**
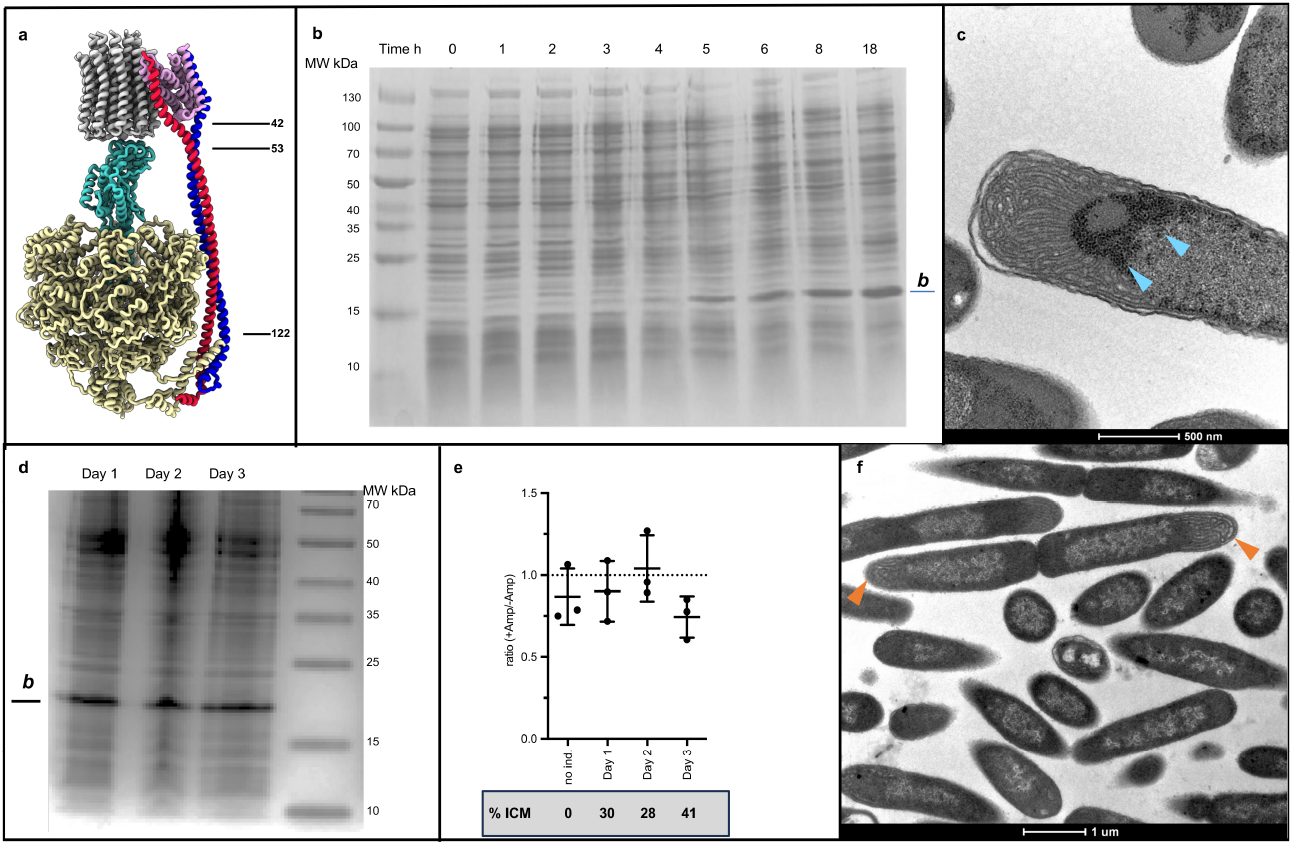
Biogenesis of subunit *b* dependent ICMs preserves cell viability. **a**, Bacterial ATP synthase structure. F_o_ domain c-ring in grey and subunit a in pink, F_1_ catalytic domain in yellow, and the central stalk in green. ATP synthase subunit *b* (red and blue) forms the peripheral stalk. **b**, Time course of expression of *E. coli* C43(DE3) cells overproducing subunit *b*. **c**, Examples of ribosome distribution (arrows) around ICMs **d**, subunit *b* levels in total cell extracts after three consecutive overnight inductions. **e**, Plasmid stability test: serial dilutions of the bacterial culture before induction and after each 18 hours induction are spread on plates with or without ampicillin. The graph shows the ratio of the number of colonies on plates on the next days. The number of ICMs containing cells is indicated in the grey box. **f**, Example of dividing cell 1 hour after dilution of the 18 hours induced culture at day 1. Orange arrows indicate ICMs

Overproduction of the entire F_1_F_0_ ATP synthase induces a low amount of ICMs[9], while overproduction of its subunit *b* alone in the T7RNA polymerase-based expression system, leads to massive ICMs accumulation[10]. The subunit *b* forms the stator and is composed of three domains: a single-span N-terminal transmembrane domain (residues 1–42) that interacts with the a subunit of the F_0_ domain, a dimerization coiled-coil domain (residues 53–122)[11], and a C-terminal domain (residues 122–156) that interacts with the *δ* subunit to form the stalk (Fig. 1, Panel a), preventing rotation of the F_1_ domain during F_0_ rotation for ATP synthesis[12].

Overexpression of the *atpF* gene, which encodes the subunit *b* of ATP synthase, is toxic to both BL21(DE3) and C41(DE3) T7 host strains. The C43(DE3) host was genetically selected to express *atpF* at high levels without toxicity[13]. In this attenuated host, production of the ATP synthase subunit *b* triggers intracellular membrane formation with a characteristic hexagonal array morphology that requires the presence of cardiolipin[10, 14].

The BL21(DE3) derivatives C41(DE3) and C43(DE3) have contributed to 18% of all membrane protein structures deposited in the Protein Data Bank (PDB)[15]. In addition, ATP synthase subunit *b*-dependent ICMs formation represents an interesting tool for *in situ* structural analysis of membrane proteins and *in vivo* encapsulation of proteins and small molecules[16]. Therefore, we aim to unravel the molecular determinants driving ICMs formation in this context.

Using omics approaches, electron microscopy, high-resolution cryo-electron tomography, and molecular dynamics simulations we show that the ATP synthase subunit *b* establishes an unexpected mode of interaction that tethers ICMs with regular spacing within the cell and that ICMs formation elicits a specific metabolic and membrane stress response pattern involving members of the ESCRT-III superfamily.

## 3 Results

### Biogenesis of subunit *b* dependent ICMs preserves cell viability

To investigate the biogenesis of ICMs, we first performed a kinetics experiment (Supplementary Fig. 1). ATP synthase subunit *b* was overexpressed in the *E. coli* C43(DE3) bacterial host for up to 18 hours post-induction at 25 °C. Four hours after induction the recombinant ATP synthase subunit *b* was detected as one of the major protein bands of *E. coli* total cell extracts (Fig. 1, Panel b). It reached its maximum protein level after overnight induction. Classical transmission electron microscopy showed distinct membrane structures over time. Six hours after induction vesicular structures became apparent (Supplementary Fig. 1, 6h), which evolved into a dense membrane network close to the pole of the cell (Supplementary Fig. 1, 18h). Up to 30% of induced cells showed dense internal membranes as previously reported [10]. In most cases ribosomes appeared to be excluded from the membrane network and accumulated around the internal membranes (Fig. 1, Panel c).

To assess the viability of the cells upon ICMs formation, a three-day-long induction experiment was set up. After the first overnight induction, the culture was diluted 1/100 times and induced again with IPTG at OD_600nm_=0.6 for 18 hours. After each overnight induction ATP synthase subunit *b* protein levels in total cell extracts remained stable, as shown in Fig. 1, Panel d. To assess plasmid stability, serial dilutions of the three overnight cultures were performed to count the number of cells able to form a colony on a plate with or without ampicillin. The ratio (+Amp/-Amp) was close to 1 (Fig. 1, Panel e) showing that cells retained the expression plasmid after three consecutive inductions. Notably, the number of intracellular membrane-positive cells, assessed by transmission electron microscopy (TEM), remained stable over the three-day experiment and reached up to 40% at day 3. (Fig. 1, Panel e). To demonstrate that intracellular membrane formation does not compromise cell division cross-sections of cells were analyzed by TEM 1 hour after dilution of the overnight culture. Figure 1 Panel f shows an example of two ICMs-positive dividing cells. We conclude that the time course of subunit *b* expression in the C43(DE3) T7 host promotes ICMs formation and maintains cell viability and division.

### Physiological adaption of bacterial cells to intracellular membranes

To assess the physiological adaptation of the cells in response to ICMs formation we performed whole-cell RNA-Seq analysis and label-free quantitative ribo-proteomics analysis on the purified ribosomal fraction. Gene and protein expression patterns were compared between cells harboring the control pHis17 expression plasmid and cells harboring the pHis17-*atpF* plasmid. Additionally, we identified the membrane proteins present in the purified intracytoplasmic membranes fraction by mass spectrometry (Supplementary Table 1).

Five minutes after the addition of IPTG, the gene expression profile of the cells (Fig. 2, Panel a) revealed up to 15 local network clusters that were significantly enriched upon expression of *atpF* (Supplementary Fig. 2, Panel a). Seven of these clusters related to the general increase of catabolism: metabolite transport of maltodextrin (*malK, lamB*), allose (*alsA*), sorbitol (*srlA*), and glycerol (*glpT*) was stimulated, as were catabolic pathways of glycerol, fatty acids (*fadL*), arginine (*astC*), allose (*alsA*), fucose (*lldD*), and the TCA cycle (*sucA, sucB*). Energy needs increased to support the expression of the recombinant *atpF* gene. The global DNA-binding transcriptional dual regulator *fis* was significantly downregulated by −2.8 log_2_. Consequently, fourteen local clusters linked to ribosome assembly and function were enriched, suggesting a moderate but global decrease in ribosome subunit gene expression.

**Fig. 2.**
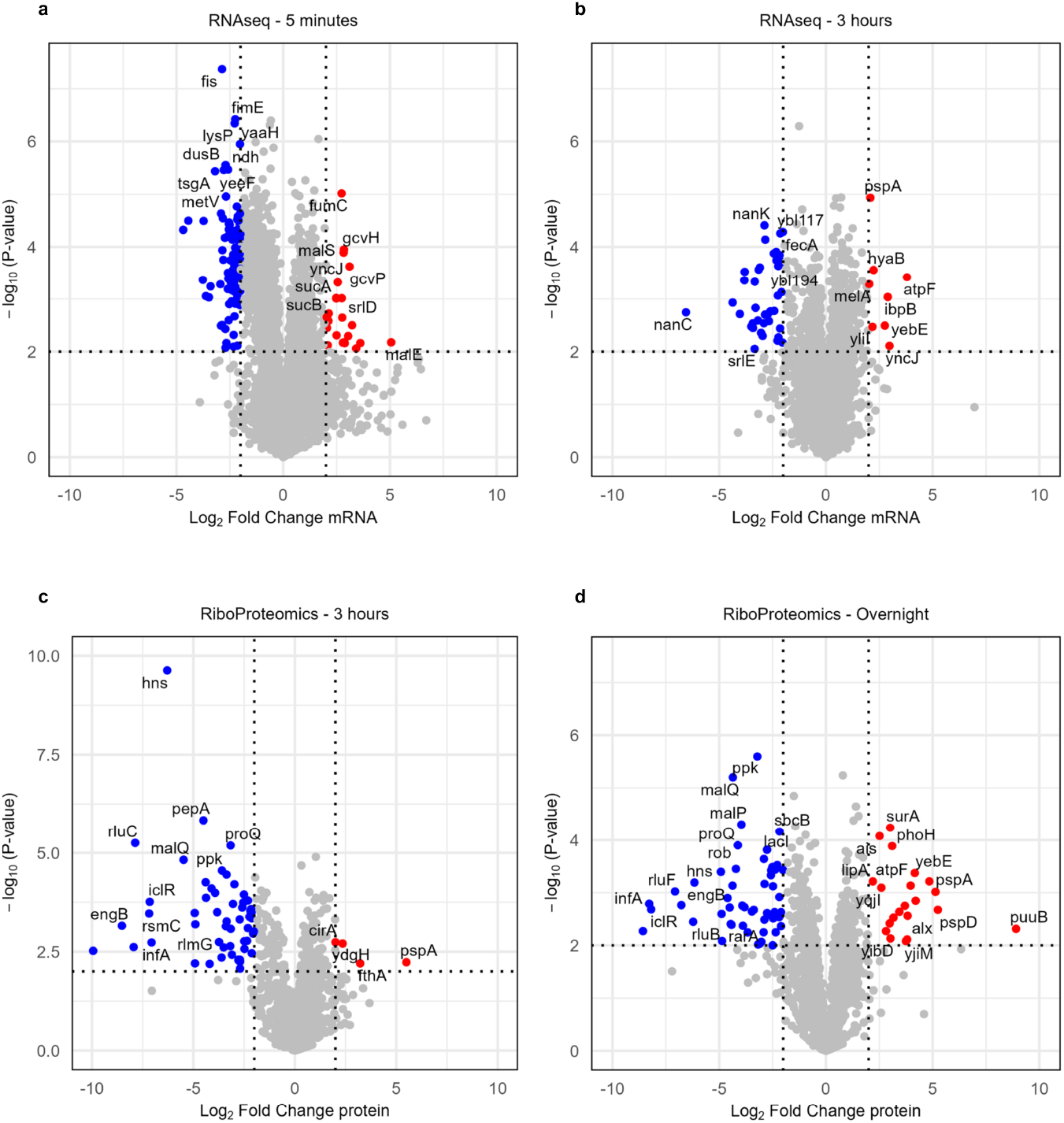
Bacterial cell response to ATP-synthase subunit *b*-dependent ICMs formation. Volcano plots showing *E. coli* C43(DE3) proteomics results for subunit *b* versus control. Data are based on *n* = 3 biological replicates for each condition. The vertical dashed lines represent a log_2_(fold change) > *±*2. Proteins above the horizontal dashed line have an adjusted *P* < 0.01. **a**, Whole cell RNAseq analysis 5 minutes after induction. **b**, Whole cell RNAseq analysis 3 hours after induction. **c**, Ribosome proteomics analysis 3 hours after induction. **d**, Ribosome proteomics analysis 18 hours after induction.

Three hours after adding IPTG, a few genes were upregulated. Besides *atpF*, transcripts for the chaperone *ibpB* and phage shock protein *pspA* were significantly increased by 2.9 log_2_ and 2.1 log_2_ respectively. In contrast, genes related to catabolic processes (galactarate, D-glutarate, aldaric acid), iron homeostasis, and metabolite transport (sorbitol, sialic acid) were decreased (Fig. 2, Panel b; Supplementary Fig. 2, Panel b). The protein unfolding and membrane permeability stress responses were also evident in the proteomic analysis of the ribosomal extract, with PspA being upregulated. Notably, all proteins from the *psp* operon, together with the membrane-bound AAA+ protease FtsH, were found in the purified intracellular membranes fraction, suggesting membrane permeability stress occurs during ICMs formation (Supplementary Table 1).

At the ribosome level (Fig. 2, Panel c), several networks of downregulated proteins involved in ribosomal RNA pseudouridine synthesis and methylation (RluB, RluC, RluD, RsmC, RlmG, RluF) negatively impacted ribosome assembly and function. DNA-binding proteins (H-NS, IhfB, IhfA, HupA), which are involved in regulating transcription and translation, were also decreased. Other upregulated local network clusters include membrane assembly, membrane invagination, and peptidase activity, with enrichment scores of 1.21, 1.24, and 1.44, respectively (Supplementary Figure 2, Panel c). This is consistent with the extensive membrane remodeling observed by electron microscopy.

After overnight accumulation of intracellular membranes, the ribosomal proteomic analysis (Fig. 2, Panel d) confirmed the general stress response observed after 3 hours with downregulation of the H-NS protein by −6.1 log_2_, upregulation of the phage shock protein pspA by 4.8 log_2_, and activation of the putrescine catabolism pathway (Supplementary Fig. 2, Panel d). To conclude, our data suggest that *E. coli* cells enter a quiescence-like state in response to both ATP synthase subunit *b* overexpression and membrane stress induced by intracellular ICMs formation.

### Cryo-electron tomography shows subunit *b* mediated regular spacing of internal membranes

To gain insight into the molecular mechanism of intracellular membrane formation high-resolution cryo-electron tomography was performed after overnight production of ATP synthase subunit *b*. Lamellae from *E. coli* cells observed by transmission electron microscopy confirmed that most bacterial cells contained ICMs (Fig. 3, Panels a and b). Several tomograms showed dark straight densities connecting intracellular membranes and to the inner membrane (Fig. 3, Panel c). Membrane segmentation as well as ribosome picking and averaging was performed on denoised tomograms (Fig. 3, Panel d, Supplementary movie 1). This revealed that the ICMs are composed of regularly organized and packed tubular membranes, mostly at the pole of the cells, delineating a ribosome exclusion zone similar to what was observed with classical TEM (Fig. 1, Panel c). Additionally, we observed that at the periphery of the ICMs membrane invaginations rooted from the inner bacterial membrane, which fed the intracellular membrane network (Fig. 3, Panel e). The dark straight densities coating the tubular membranes were overall normal to the membrane surface and defined an average 26 nm distance between the membrane tubes. We hypothesized that the dark densities were ATP synthase subunit *b*. However, considering the small size of this molecule (a single coiled-coil domain) it was not possible to pick them and perform subtomogram averaging.

**Fig. 3.**
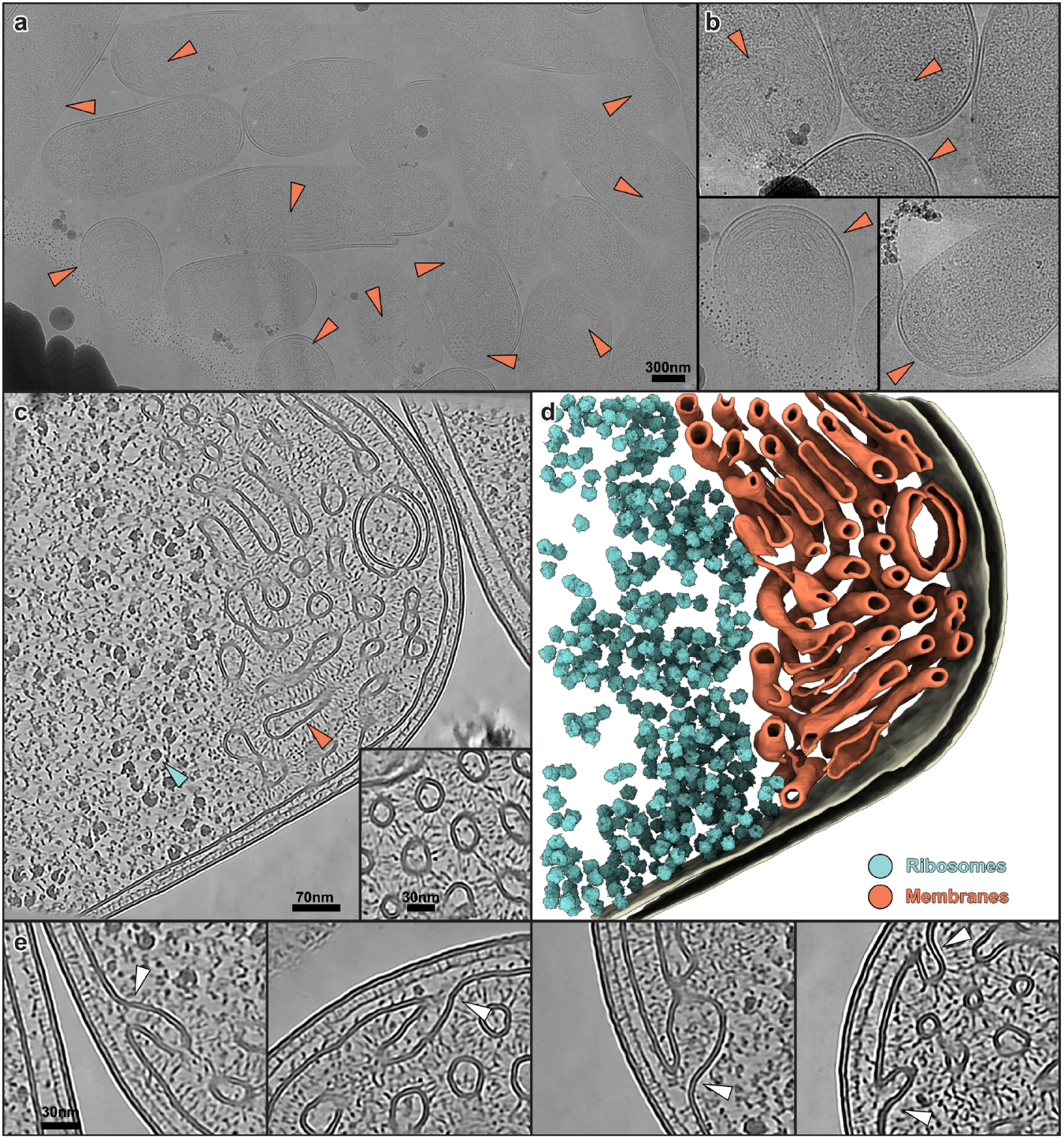
Cryo-electron tomography analysis of *E. coli* C43(DE3) upon ATP synthase subunit *b* overexpression. **a**, Transmission electron microscope overview of a lamella from subunit *b* expressing *E. coli* cells. Orange arrows indicate the presence of a membrane network. **b**, Zoomed-in snapshots of areas with membrane networks. **c**, Slice of a tomogram depicting a bacterial cell with the membrane network. Arrowheads point to ribosomes (blue) and membranes (orange). The insert shows a zoomed-in view of the membranes. **d**, Corresponding segmentation of the tomogram with single-particle analysis (STA) structure of ribosomes mapped back into their respective particle positions (blue). Outer and inner bacterial membranes are shown in beige, and the internal membrane network in orange. **e**, Zoomed-in views from other tomograms showing that the membrane network originates from the plasma membrane. Junctions are indicated with white arrows.

### ATP synthase subunit *b* C-terminal domain is required for ICMs accumulation

Next, we aimed to discern the molecular interactions responsible for the formation of ICMs. We dissected the ATP synthase subunit *b* protein into two deletion variants: b(1–120) comprising the *α*-helix (residues 63–120) that forms a coiled-coil interaction between two ATP synthase subunit *b* monomers, and b(1–87), retaining only half of the previously mentioned helix (Table 1). Following overnight induction at 25 °C the harvested cells were processed and subjected to TEM analysis (Fig. 4, Panel a). Both ATP synthase subunit *b* deletion constructs exhibited distinct ICMs phenotypes compared to the wild-type ATP synthase subunit *b*. Additionally, the proportion of ICMs-containing cells decreased from 30% to 6% and the remaining intracellular membranes appeared disordered and lacked interconnections. Hexagonal tubular structures connected by dark, straight densities were not found in any of the subunit *b* deletion variants (Fig. 4, Panel a).

**Table 1.**
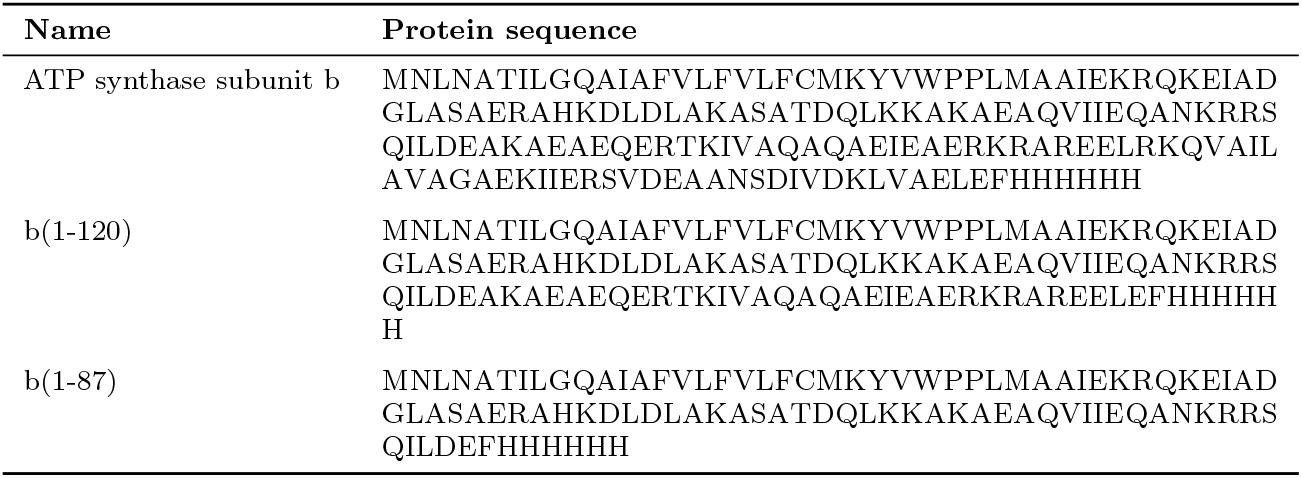
Protein sequences of ATP synthase subunit b constructs.

**Fig. 4.**
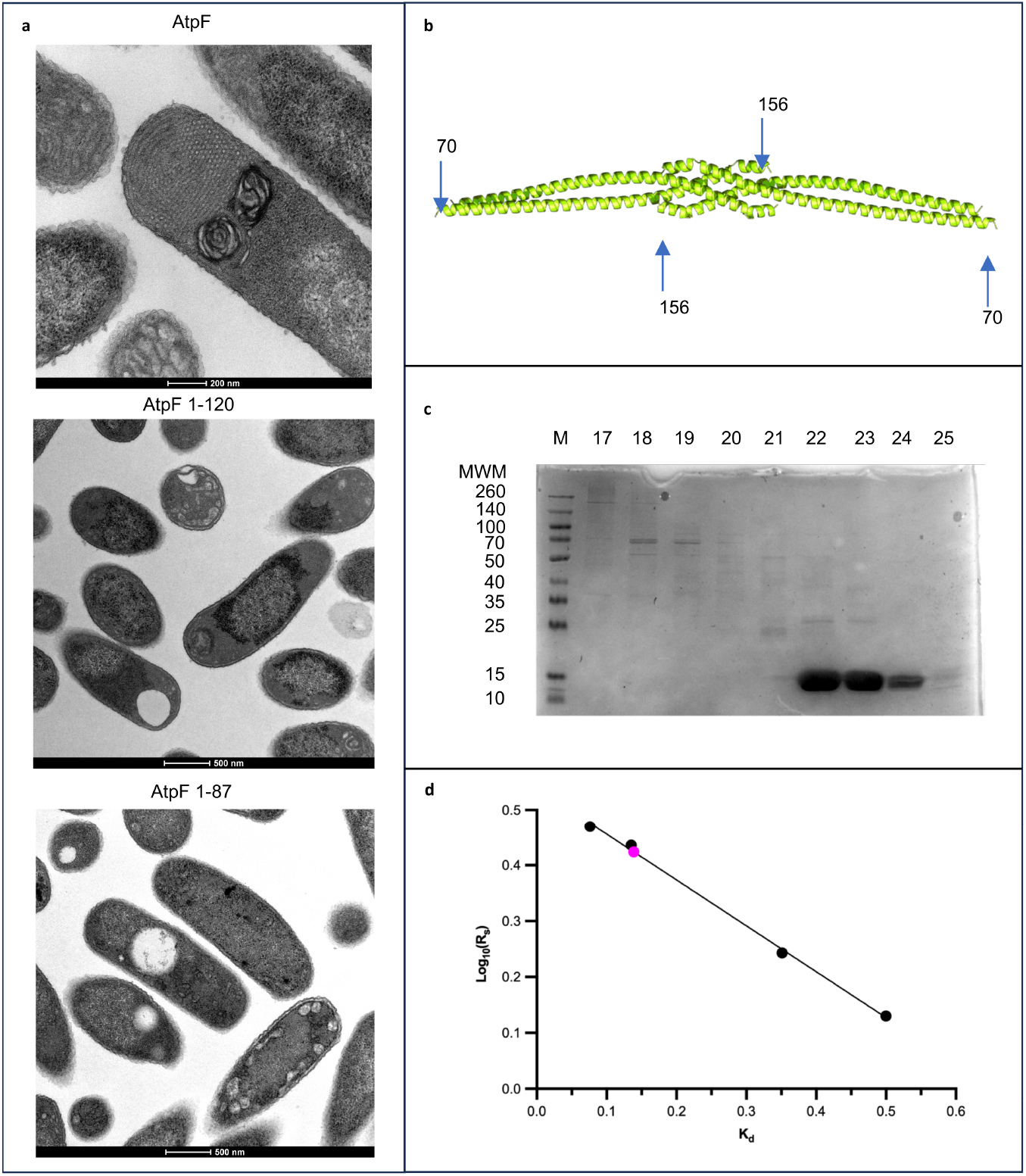
The C-terminal domain of the subunit *b* is essential for ICMs formation. **a**, Transmission electron microscopy images showing the intracellular morphologies upon overproduction of ATP synthase subunit *b* deletions. **b**, AlphaFold3 model of the C-terminal 86 amino acids of *b*(70–156). **c**, Coomassie blue-stained SDS-PAGE of purified *b*(70–156). **d**, Size estimation of purified *b*(70–156), shown by a magenta dot, by gel filtration chromatography. The black dots represent the calibration standards.

We hypothesized that the C-terminal domain of ATP synthase subunit *b* (residues 120–156) mediates the regular spacing between ICMs. AlphaFold predicts the structure of the full-length ATP synthase subunit *b* as observed in the F_o_F_1_ ATP synthase complex (PDB: 6OQT) [17]. In contrast, the subunit *b*(70-156) and *b*(90-156) domains are predicted as a tetramer with head-to-head interactions at the C-terminal extremities. However, given the low confidence score of the predictions, the stability of the truncated subunit *b* tetramers (Fig. 4, Panel b and Fig. 5, Panel a) was tested both experimentally and by molecular dynamics simulations.

**Fig. 5.**
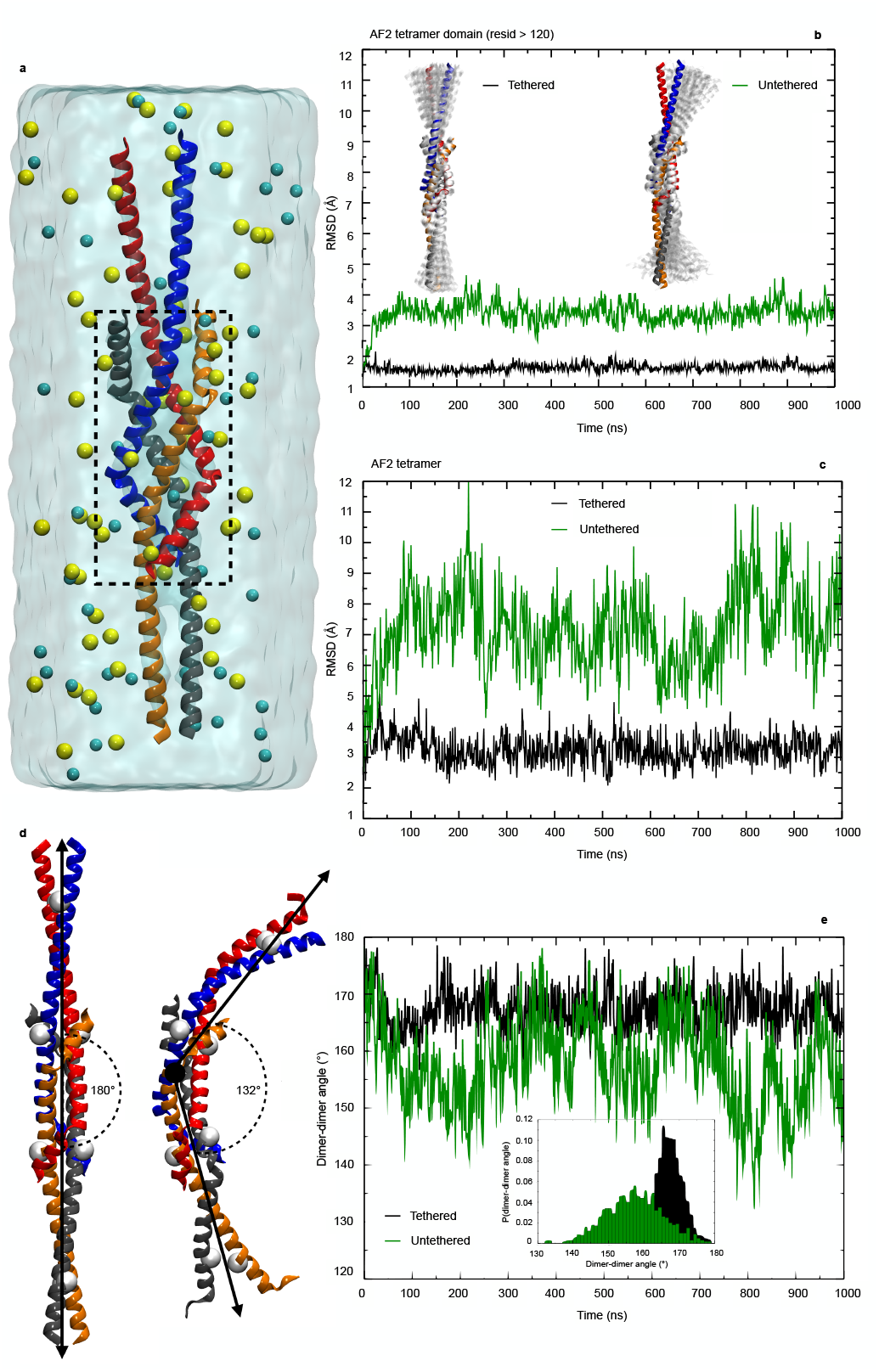
Structural dynamics of an AlphaFold2-predicted tetramer of the *E. coli* ATP synthase subunit *b* (residues 90–156). **a**, Molecular dynamics (MD) simulation setup of the AlphaFold2 tetramer solvated in an explicit water box (light blue transparent surface) containing 0.150 M NaCl (Na^+^ and Cl^-^ ions shown as yellow and cyan spheres, respectively). The dashed rectangle highlights the tetramerization domain (residues 120–156). **b**, Time evolution over 1 µs MD trajectories of the backbone root-mean-square deviations (RMSD) for the tetramerization domain. Black curves correspond to a trajectory in which the tetramer is tethered (the four terminal Ala90 residues are harmonically restrained to their initial positions), whereas green curves correspond to a trajectory in which the tetramer is untethered. Insets show 50 conformations (transparent white cartoons) sampled along the MD trajectories for the tethered (left) and untethered (right) tetramers, overlaid on the initial AlphaFold2 model (solid-colored cartoons). **c**, Time evolution of the RMSD for the full AlphaFold2 tetramer. **d**, Tetramer structures representing the extreme angles between the two forming dimers, sampled along MD trajectories. Arrows indicate the axes used to measure the angle, and white spheres mark the C*α* atoms defining these axes. **e**, Time evolution of the angles formed by the two dimers for tethered (black line) and untethered (green line) tetramers. The inset shows the probability distribution functions of these angles.

Subunit *b* tetramer *b*(70–156) was produced in C43(DE3) cells, purified, and subjected to gel filtration on a Superdex75 column (Fig. 4, Panel c). Based on column calibration in the same buffer (Fig. 4, Panel d), an apparent molecular weight of 47.6 kDa was calculated [18]. This apparent molecular weight was consistent with the 43.4 kDa molecular weight of a tetramer of the subunit *b*(70-156).

Next, we investigated the stability and dynamics of the head-to-head interactions in subunit *b*(90-156) by molecular dynamics (MD) simulation (Fig. 5, Panel a and Supplementary Fig. 3). Stability was tested with or without anchoring the N- and C-terminal extremities. RMSD analysis of subunit *b*(90-156) showed that the tetrameric complex was maintained over 1 *µ*s MD simulation with high flexibility of the *b*(90-100) dimeric region (Fig. 5, Panels b and c). We then assessed the consequence of this flexibility on the full-length subunit *b* tetramer. Both extreme conformations (180° and 132°) are shown in Fig. 5, Panel d. The distribution of angles between both vectors (100-156) of the tetramer is shown in Fig. 5, Panel e. We conclude that the C-terminal domain of subunit *b* mediates unusual head-to-head interaction, which could explain the dark straight densities observed by cryo-tomography connecting intracellular membranes and to the inner membrane.

### Membrane tethering by ATP synthase subunit *b*

To assess this hypothesis, we calculated the theoretical distribution of membrane-to-membrane distances mediated by subunit *b*. First, the full-length ATP synthase subunit *b* was modeled in a lipid bilayer. The transmembrane span is 42 amino acids long (Supplementary Fig. 4, Panel a), consistent with protease analysis of subunit *b* topology. Second, based on the topology of subunit *b*, a model of subunit *b*-mediated tethering of membranes is shown in Fig. 6, Panel a, in which membrane-to-membrane distances are inferred from the Asp42-Asp42 (green spheres) (see Supplementary Fig. 4, Panel b). To experimentally validate the model, distance measurements from membrane-to-membrane (Fig. 6, Panel b) on 6 different tomograms showed an average spacing between internal membrane structures of 26 nm (Fig. 6, Panel c). Membrane-to-membrane distance was also calculated based on the distribution of angles in the subunit *b* tetramers (Fig. 5, Panel e). The calculated mean value of 26 nm (Fig. 6, Panel c) equaled the experimental value, and the distribution of membrane-to-membrane distances inferred from the model predicted 75% of the experimentally measured distances. We conclude that the model of membrane tethering mediated by subunit *b* explains the final organization of ICMs accurately.

**Fig. 6.**
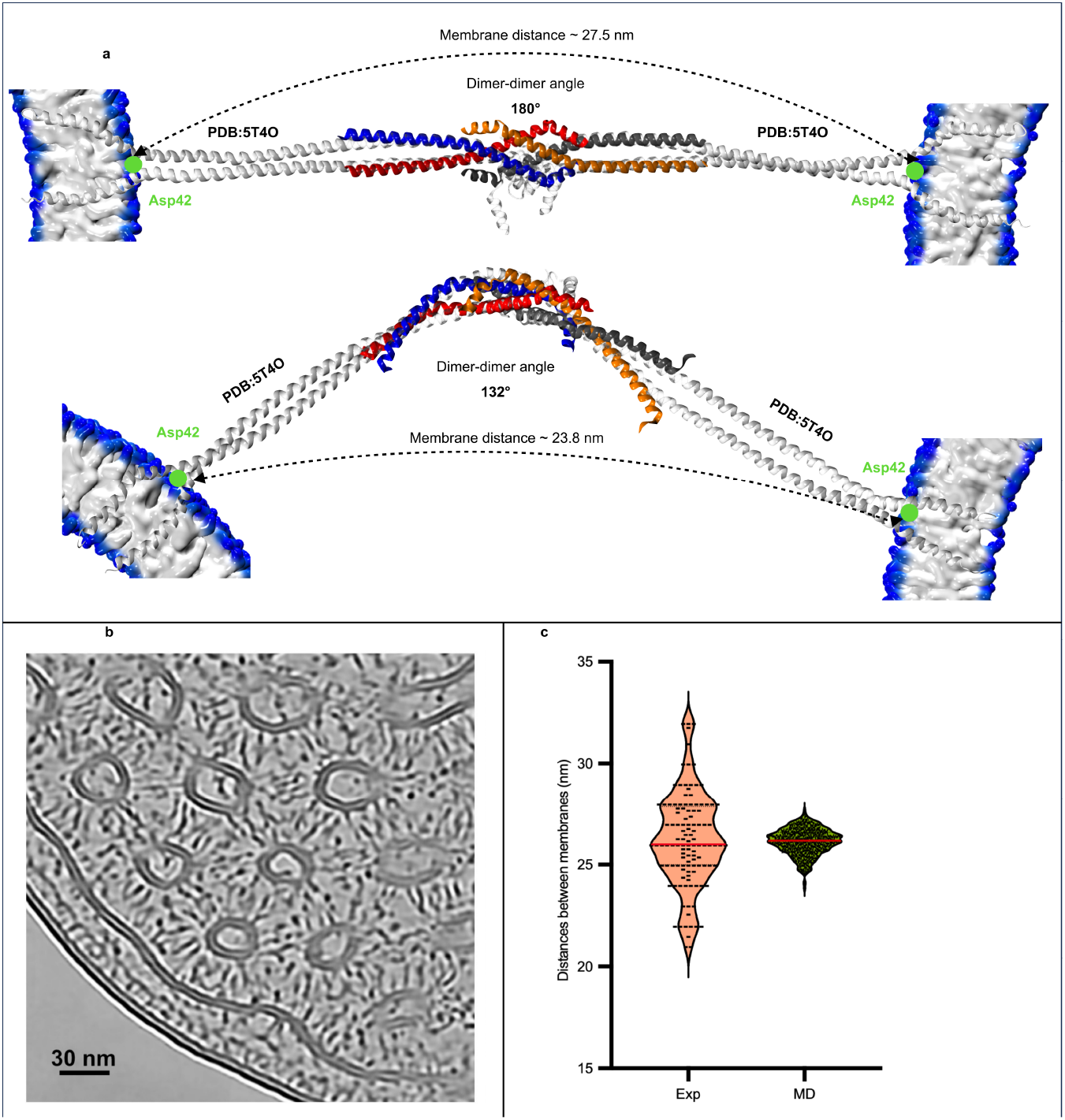
Comparison of membrane-to-membrane distances measured from cryo-electron tomograms and predicted by molecular modeling. **a**, Illustration of the modeling strategy employed to infer membrane-to-membrane distances for AlphaFold2 (AF2) tetramer structures featuring two extreme angles between their two forming dimers sampled by molecular dynamics (MD). For each conformation of the untethered tetramer sampled along a 1 µs-long MD trajectory, two copies of the full-length *E. coli* ATP synthase subunit *b* dimer (PDB: 5T4O[50]) (white cartoons) are aligned on each dimer of the AF2 tetramer (solid-colored cartoons). Theoretical membrane-to-membrane distances are inferred from the Asp42-Asp42 distances (green spheres) (see Supplementary Fig. 3). **b**, Zoomed-in view of a tomogram slice from which membrane-to-membrane distances within the membrane network of ATP synthase subunit *b*-expressing *E. coli* cells are measured. Scale bar: 50 nm. **c**, Violin plots of the membrane-to-membrane distance distributions measured experimentally (left) and inferred from molecular modeling (right). The dashed line indicates the mean experimental distance of 26 nm.

## 4 Discussion

Our results provide fundamental new insights into how the ATP synthase subunit *b* can induce distinct membrane microcompartments within the bacterial cell. Our findings reveal the different steps and mechanisms involved in ICMs formation. First, ICMs initiate with the appearance of a large vesicle within the cytoplasm; second, ICMs accumulate at the pole through invagination of the inner membrane; third, intracellular membrane structures exhibit an average spacing of 26 nm, mediated by head-to-head interactions of subunit *b* tetramer; fourth, bacterial cells enter into reversible metabolic quiescence. The gene and protein expression patterns observed here reveal the physiological adaptations of the cell to subunit *b* production and ICMs formation. Five minutes after induction whole-cell transcriptomic analysis shows an increase in energy demand, which is exemplified by the overexpression of metabolite transporters. Three hours after induction of subunit *b* cells enter quiescence, downregulate the translation machinery, and exhibit membrane remodeling and protein folding stress. Notably, lipid metabolism genes are not upregulated showing that ICMs accumulation does not require increased activity of the lipid biosynthesis machinery. This observation aligns with proteome analyses by Gubellini *et al*.[19], who reported no stimulation of lipid biosynthesis upon overproduction of five different membrane proteins. In addition none of the major changes in gene expression patterns (such as sigma factors, proteases, flagella, acid resistance, or protein translocation) observed by Gubellini and coworkers—or in response to cell stress induced by heat, cold, oxidative stress, nitrogen starvation, or antibiotic treatment[4]—are found with ATP synthase subunit *b* overproduction. In contrast, one of the most significant changes observed by whole-cell RNAseq and riboproteomics analyses is the upregulation of the phage shock protein PspA. Of note, PspA protein is also identified by MS in purified subunit *b* containing ICMs. The PspA protein could increase membrane curvature at the pole of the inner membrane, bind cardiolipin, and participate in ICMs formation, as proposed by Hudina and coworkers[20]. We previously showed that subunit *b*-dependent ICMs are enriched in cardiolipin and deletion of the three *cls A, B* and *C* genes alters ICMs morphology into onion-like structures [14]. We propose the following mechanism for ICMs cardiolipin enrichment. ICMs accumulate during the late stationary phase at the bacterial cell poles, which contain cardiolipin-enriched inner membranes [21]. Cryo-EM images (Fig. 3, panel e) show direct connections between ICMs and invaginating inner membranes, resulting in cardiolipin transfer to ICMs.

The model of ICMs organization during the late stationary phase, presented in Figure 6, Panel a, is supported by several lines of evidence. First, high-resolution cryo-electron tomography shows that the ICMs are sequestered within the cytoplasm by a dense network of protein arrays bridging membranes and vesicles. Second, the *b*(120–156) C-terminal domain is mandatory for ICMs formation. Third, AlphaFold predictions combined with molecular dynamics simulations of the subunit *b* C-terminal domain and biochemical analyses reveal a tetrameric assembly with head-to-head interactions bridging two separate membrane structures. Fourth, the model inferred from these data accounts for 75% of the membrane-to-membrane distance distribution measured from cryo-EM tomograms. In eukaryotic cells the dimerization of F_1_F_o_ ATP synthase plays a critical role in shaping the inner mitochondrial membrane [22]. Specifically, the membrane-embedded domains of the peripheral stalk subunits (e.g., subunits e and g in mammals) form the dimerization interface. Structural analyses reveal that the angle between the rotary axes of the monomeric complexes spans a broad range (0°–95°), reflecting distinct dimer conformations across species and tissues [22]. These dimers, and their higher-order oligomers, actively induce local membrane curvature [23] and are essential for the characteristic folding of mitochondrial cristae [24, 25]. In the subunit *b*-dependent membrane network the transmembrane span of the *E. coli* subunit *b* does not play a significant role. Instead, we demonstrate that the C-terminal soluble domain of subunit *b* forms supramolecular membrane structures through head-to-head interactions. Whether this property has been conserved in other prokaryotes to generate F_1_F_o_ ATP synthase-enriched microcompartments or in eukaryotes to shape energy-transducing membranes, remains to be investigated.

The proposed mechanism of protein-protein interactions bridging membranes may explain other examples of ICMs organization in *E. coli*, which exhibit regular spacing between membrane structures. Such examples include ICM formation upon fumarate overproduction [26], glycosyltransferase LpxB overexpression [27], and Tsr receptor clustering [28]. In conclusion, our results elucidate the molecular mechanism underlying the membrane-remodeling function of the ATP synthase subunit *b*. This represents a critical advance toward the design of synthetic membrane organelles in *E. coli* for the *in vivo* structural analysis of heterologously produced membrane proteins.

## 5 Material and Methods

### *E. coli* strain and culture conditions

The *E. coli* ATP synthase subunit *b* gene *atpF* was inserted into the pHis17 T7 expression vector, a derivative of the pMW7 plasmid[13, 29]. In pHis17 the multiple cloning site (MCS) was modified by inserting the following sequence CATATGGGATCCC ATCATCATCATCATCATTAAAAGCTTCACCACCACCACCACCACTAAGAATTCCATCATCATCATCATCA TTAA between the *NdeI* and *EcoRI* sites. As a result a C-terminal 6xHis tag is added in-frame downstream of the *BamHI, HindIII*, and *EcoRI* sites of the polylinker. The *atpF* gene was subcloned between the *NdeI* and *EcoRI* restriction sites, resulting in an EFHHHHHH C-terminal tag.

Transformed bacteria were selected on 2-YT agar plates supplemented with ampicillin (100 µg mL^-1^). For protein expression individual recombinant colonies were inoculated into 100 mL of liquid 2-YT medium containing ampicillin (100 µg mL^-1^) and cultured at 37 °C until the OD_600_ reached 0.4. The cultures were then induced by adding 0.5 mM IPTG and the temperature was reduced to 25 °C for overnight protein expression.

### Transmission Electron Microscopy

Samples overexpressing the proteins of interest were fixed with 2.5% glutaraldehyde in 0.1 M sodium cacodylate buffer overnight at 4 °C, postfixed in 1% osmium tetroxide + 1% potassium ferrocyanide for 2 hours at 4 °C, dehydrated in ethanol, and infiltrated in a mixture of EMbed 812 (Electron Microscopy Sciences) epoxy resin and propylene oxide (1:1) overnight. The samples were then embedded. Ultrathin sections were obtained with a Reichert-Yung Ultracut ultramicrotome, collected on 200 mesh copper grids, and counterstained with uranyl acetate and lead citrate. Samples were examined with a Tecnai G2 (FEI) transmission electron microscope operating at 120 kV, and digital images were acquired using a Veleta (Olympus Soft Imaging Solutions) digital camera.

### RNA Extraction and Sequencing

Total RNA from *E. coli* was extracted using a modified phenol-chloroform protocol [30]. Briefly, 700 µl of cell culture were harvested and resuspended in a lysis solution containing 35.5 µl of 20% sodium dodecyl sulfate (SDS), 7 µl of 200 mM Na-EDTA (pH 8.0), and 500 µl of water-saturated phenol (preheated to 65°C). The mixture was vigorously vortexed for 30 seconds and incubated at 65°C for 5 minutes with intermittent mixing. After cooling on ice for 2 minutes, the samples were centrifuged at 16,000 × g for 2 minutes at 4°C to separate the aqueous and organic phases. The aqueous phase was carefully transferred to a new RNase-free microcentrifuge tube, and the extraction was repeated twice more: first with an equal volume of water-saturated phenol, and then with a 1:1 mixture of phenol:chloroform (24:1 chloroform:isoamyl alcohol). RNA was precipitated by adding 1/10 volume of 3 M sodium acetate (pH 5.2) and 2.5 volumes of ice-cold 100% ethanol, followed by incubation at -20°C for 2 hours. The RNA pellet was collected by centrifugation at 16,000 × g for 20 minutes at 4°C, washed once with 1 ml of 70% ethanol, air-dried for 10 minutes, and resuspended in 400 µl of RNase-free sterile water.

To eliminate residual genomic DNA the RNA samples were treated with RNase-free DNase I (Qiagen) at 37°C for 30 minutes according to the manufacturer’s instructions. A second round of phenol:chloroform extraction and ethanol precipitation was then performed to ensure RNA purity. The final RNA pellet was resuspended in 50 µl of RNase-free water. RNA concentration and purity were assessed using a NanoDrop ND-1000 spectrophotometer (Thermo Fisher Scientific). Purity was evaluated by the A_260_/A_280_ ratio, with values between 1.8 and 2.1 considered acceptable. RNA integrity was verified by depositing 1 µg of each sample on a 0.7% (w/v) agarose gel stained with ethidium bromide, and visualizing the 16S and 23S rRNA bands under UV light.

RNA sequencing was performed at the Transcriptome and Epigenome Platform of the Institut Pasteur (Paris, France). RNA quality and quantity were confirmed using a Bioanalyzer 2100 (Agilent Technologies). For library preparation, ribosomal RNA (rRNA) was depleted from 10 µg of total RNA using the Ribo-Zero rRNA Removal Kit (Epicentre, Singapore) according to the manufacturer’s protocol. Non-directional cDNA libraries were prepared from enriched, fragmented mRNA using the TruSeq Stranded RNA LT Sample Prep Kit (Illumina, sets A and B). cDNA fragments of approximately 150 base pairs (bp) were ligated with Illumina adapters, PCR-amplified, and purified from each library.

Sequencing was performed on an Illumina HiSeq 2000 platform, generating single-end reads of 51 bases. Raw sequencing reads were aligned to the *E. coli* BL21(DE3) reference genome (NC 012892.2) using Bowtie2 (version 2.2.5) [31]. Differential gene expression analysis was performed using the DESeq2 package (version 1.8.2) [32] in R (version 3.2.0) with Bioconductor [33]. Normalization of read counts and differential expression testing were carried out according to the DESeq2 model. To control the false discovery rate (FDR), a Benjamini-Hochberg (BH) p-value adjustment was applied, with a significance threshold set at FDR *<* 0.05.

### Purification of *E. coli* ribosomes

Highly purified ribosomes were isolated from C43(DE3) cells expressing either the empty pHis17 plasmid (control) or pHis17 containing the *atpF* gene. The preparation was performed via sucrose cushion centrifugation, modified from a previous protocol[34]. One-liter cultures of both control and ATP synthase subunit *b* over-expressing cells, incubated overnight at 25 °C, were harvested by centrifugation at 4,500 × *g* for 20 minutes at 4 °C. Cells were then washed in Buffer A (20 mM Tris pH 7.4, 10 mM MgOAc, 100 mM NH_4_OAc, 0.5 mM EDTA). All subsequent steps were performed at 4 °C. The cell pellet was resuspended in Buffer A supplemented with 0.1 mg mL^-1^ lysozyme, 6 mM *β*-mercaptoethanol, and 0.1% (v/v) protease inhibitor cocktail. Cells were lysed by two passages through a French pressure cell at 10,000 psi.

The lysate was clarified by two sequential centrifugations at 22,000 × *g* for 15 minutes at 4 °C. The resulting supernatant was gently overlaid onto an equal volume of 37.7% (w/v) sucrose cushion prepared in Buffer B (20 mM Tris pH 7.4, 10 mM MgOAc, 500 mM NH_4_OAc, 0.5 mM EDTA, 6 mM *β*-mercaptoethanol). Ribosomes were pelleted by ultracentrifugation in a Beckman Coulter Type 70 Ti rotor at 144,000 × *g* for 20 hours at 4 °C. Following decanting of the sucrose cushion, the ribosomal pellet was resuspended in Buffer C (20 mM Tris pH 7.4, 7.5 mM MgOAc, 60 mM NH_4_OAc, 0.5 mM EDTA, 6 mM *β*-mercaptoethanol). Purified ribosomes were quantified using a NanoDrop. Each condition for control and ATP synthase subunit *b* (3 hours and overnight) had three biological replicates, each analyzed by SDS-PAGE and Coomassie staining. The gel was further used for proteomic analysis using label-free quantification.

### Mass spectrometry

In-gel digestion: after a 1 cm short migration of 10 µg of each protein sample by SDS-PAGE, proteins were fixed and stained with Coomassie Brilliant Blue R-250. Lanes containing proteins were excised manually and subjected to manual in-gel digestion with modified porcine trypsin (Trypsin Gold, Promega). Briefly, after destaining, bands were subjected to a 30-minute reduction step at 56 °C in the dark with 10 mM dithiothreitol in 50 mM ammonium bicarbonate (AMBIC), followed by a 1-hour cysteine alkylation step at room temperature in the dark with 50 mM iodoacetamide in 50 mM AMBIC. After dehydration under vacuum, bands were re-swollen with 250 ng of trypsin in 200 µL 50 mM AMBIC, and proteins were digested overnight at 37 °C. Supernatants were kept, and peptides present in gel pieces were extracted with 1% (v/v) trifluoroacetic acid and dried in a vacuum concentrator. Peptides were then solubilized in 50 µL of solvent A (0.1% (v/v) formic acid) to a final concentration of 200 ng µL^-1^.

Mass spectrometry analyses were performed on a Q-Exactive Plus hybrid quadrupole-orbitrap mass spectrometer (Thermo Fisher, San José, CA, USA) coupled to a Neo Vanquish liquid nano-chromatography system (Thermo Scientific). Peptide mixtures were analyzed in duplicate. Five microliters of peptide mixtures were loaded onto a Neo pepmap trap column (300 µm *×* 5 mm, 5 µm, 100 Å; Thermo Fisher Scientific) equilibrated in solvent A (0.1% formic acid) and separated at a constant flow rate of 300 nL min^-1^ on a PepMap NEO™ RSLC C18 Easy-Spray column (75 µm *×* 50 cm, 2 µm, 100 Å; Thermo Scientific) with a 90-minute gradient (3 to 20% B solvent (80% ACN, 0.1% formic acid (v/v)) in 60 minutes and 20 to 35% B solvent in 30 minutes).

Data acquisition was performed in positive and data-dependent modes. Full scan MS spectra (mass range m/z 400–1600) were acquired in profile mode with a resolution of 70,000 (at m/z 200), and MS/MS spectra were acquired in centroid mode at a resolution of 17,500 (at m/z 200). All other parameters were kept as described previously[35].

Data processing and label-free quantification: raw data were processed using the MaxQuant software package (version 1.6.0.13)[36]. Protein identifications and target-decoy searches were performed using the Andromeda search engine and a SwissProt *Escherichia coli* K12 strain database (21/11/2023; 4530 entries) in combination with the MaxQuant contaminants database (245 contaminants). The mass tolerance in MS and MS/MS was set to 10 ppm and 20 mDa, respectively. Methionine oxidation and protein N-terminal acetylation were considered as variable modifications, whereas cysteine carbamidomethylation was considered as a fixed modification. Trypsin was selected as the cutting enzyme, and 2 missed cleavages were allowed. Proteins were validated if at least ≥ 2 unique peptides with a protein FDR ¡ 0.01 were identified. The setting “Match between runs” was also used to increase the number of identified peptides. For quantification, unique and razor peptides with a minimum ratio count ≥ 2 unique peptides were used. Protein intensities were calculated by Delayed Normalization and Maximal Peptide Ratio Extraction (MaxLFQ)[37].

Statistical analysis was performed using Perseus software (version 1.6.15.0)[36]. Proteins belonging to contaminants and decoy databases were filtered. For each biological replicate the median intensity of the two injected replicates was determined, and proteins having quantitative data in at least 3 biological replicates were considered for statistical analysis using a Benjamini-Hochberg test with a threshold p-value *<* 0.05. Proteins exhibiting a Log_2_(fold change) ≥ 2 and a false discovery rate (FDR)*<* 0.01 were classified as significantly differentially expressed between the two conditions.

### Cryo-electron tomography acquisition and processing

A 4.5 µL cell suspension, grown in 2-YT media induced for 18 hours in the presence of 0.5 mM IPTG, was applied to R2/1 copper grids (Quantifoil) and plunged frozen in liquid ethane by back-sided blotting using a Leica EM GP2 (4 s blot time and 90% humidity).

First, cells were thinned down by FIB milling as described previously[38] using an Aquilos 2 microscope (Thermo Fisher Scientific). Then, lamellae were transferred to a Titan Krios G4 transmission electron microscope operated at 300 kV, equipped with a SelectrisX energy filter (slit set to 10 eV) and a Falcon 4i direct electron detector (Thermo Fisher Scientific). Dose-fractionated movies were recorded in TIFF format. The microscope was set to a nominal magnification of 64,000*×*, corresponding to a pixel size of 1.98 Å at the sample level. Dose-symmetric tilt series were recorded using the TEM Tomography 5 software (Thermo Fisher Scientific) in a tilt span of *±*60°, covered by 2° steps starting at either *±*10° offset to compensate for the lamella pretilt. Target defocus was set for each tilt-series in a range of -2 µm to -4 µm in steps of 0.5 µm, using a dose of 2.1 e^-^/Å^2^ per tilt image.

Tomograms were processed using Scipion[39] following the workflow described previously[40]. Briefly, raw frames were motion corrected using MotionCor2[41], and images were aligned with AreTomo[42] after manual tilt-series curation. Tomograms were dose filtered and reconstructed using IMOD[43]. For visualization, tomograms were denoised from their reconstructed half-volumes using Icecream[44]. Membrane segmentation was performed using MemBrain v2[45].

Ribosome identification was performed on CTF-corrected tomograms by template matching using pytom-match-pick[46]. 3D classification and averaging of ribosomes were performed in RELION[47]. Visualization and mapping back of the particles were done in ChimeraX[48] using the ArtiaX plugin[49].

### Purification of ATP synthase *b*(70–156)

ATP synthase *b*(70–156) was produced in C43(DE3) cells as for the whole ATP synthase subunit *b*, using 0.5 mM IPTG and overnight induction at 25 °C. The cells (0.5 L) were lysed using a cell disruptor at a pressure of 2 kBar (Constant Systems Ltd) in buffer A containing 50 mM phosphate (pH 7.5) and 300 mM NaCl. Cell debris was removed by centrifugation at 2,500 × *g* for 10 minutes. Aggregates were removed at 10,000 × *g* for 20 minutes. The soluble fraction was recovered after centrifugation of the 10,000 × *g* supernatant at 100,000 × *g* for 1 hour at 4 °C.

ATP synthase *b*(70–156) was purified on Ni-NTA resin (Thermo Fisher Scientific) in batch mode. The 30 mL 100,000 × *g* supernatant was incubated with 1 mL Ni-NTA resin for 1 hour and then loaded onto a 10 mL column (Bio-Rad). After removal of the breakthrough, the column was washed with 10 mL buffer A and 10 mL buffer A supplemented with 20 mM imidazole. The ATP synthase *b*(70–156) was eluted with 300 mM imidazole in buffer A. Two fractions of 0.5 mL were pooled and loaded onto a PD10 column equilibrated with buffer A. Since ATP synthase *b*(70–156) does not contain aromatic residues, protein concentration was estimated at 4 mg mL^-1^ by Coomassie-stained SDS-PAGE using BSA as a standard.

### Analytical Gel Filtration

Purified ATP synthase *b*(70–156) (100 µL, 0.8 mg) was applied to a Superdex 75 Increase 10/300 GL column and eluted at a flow rate of 0.8 mL min^-1^ in 10 mM phosphate (pH 7.5), 140 mM NaCl, and 1 mM DTT. The column was calibrated using the following standards (molecular weight, Stokes radius): conalbumin (75 kDa, 2.95 nm), ovalbumin (43 kDa, 2.73 nm), ribonuclease A (13.7 kDa, 1.75 nm), and aprotinin (6.5 kDa, 1.35 nm). The elution volume and Stokes radius of the standards were used to establish a calibration curve. The Stokes radius of the ATP synthase *b*(70–156) construct was determined from this calibration curve. Thereafter, the Stokes radius was plotted as a function of molecular weight for both protein standards and the ATP synthase *b*(70–156) construct.

### Molecular simulation details

The putative structures of the ATP synthase *b*(90–156) tetramer were generated using AlphaFold 2 and AlphaFold 3 (see Fig. 5, Panel a and Supplementary Fig. 3). The predicted assemblies correspond to tetramers composed of two dimers arranged in a head-to-head configuration. For each construct, the model with the highest ranking according to internal confidence metrics (pLDDT and predicted aligned error) was selected for subsequent simulations. Each tetramer was placed in a rectangular simulation box (56 × 58 × 163 Å^3^) and solvated with 14,742 explicit water molecules. Sodium and chloride ions were added to neutralize the system and achieve a final salt concentration of 150 mM NaCl. The full-length ATP synthase *b* dimer was extracted from the autoinhibited structure of the *Escherichia coli* ATP synthase [50] (PDB ID: 5T40), available in the Protein Data Bank. The dimer was embedded in a heterogeneous lipid bilayer composed of 70 % POPE (1-palmitoyl-2-oleoyl-sn-glycero-3-phosphoethanolamine), 15 % POPG (1-palmitoyl-2-oleoyl-sn-glycero-3-phosphoglycerol), 15 % tetrapalmitoyl cardiolipin (1^*′*^,3^*′*^-bis(sn-1,2-dipalmitoyl-glycero-3-phospho)-sn-glycerol) and fully solvated with explicit water molecules using CHARMM-GUI [51, 52]. The final molecular system consisted of a simulation box 84 × 84 × 229 Å^3^ containing 140 POPE, 30 POPG, 30 cardiolipins, 43,526 water molecules with 150 mM NaCl.

The CHARMM36 force field [53, 54] was used to describe proteins, ions and lipids, and water molecules were modeled using TIP3P [55]. All simulations were performed using NAMD version 3 [56]. Temperature was maintained at 300K using the stochastic velocity rescaling method [57] and pressure was maintained at 1 atm using the Langevin piston method [58]. A cutoff of 12 Å was applied for short-range van der Waals interactions, with a switching function starting at 10 Å. Long-range electrostatic interactions were computed using the Particle Mesh Ewald (PME) method [59]. The SHAKE/RATTLE algorithms [60, 61] were used to constrain covalent bonds involving hydrogen atoms and the SETTLE algorithm [62] was used to constrain water geometry.

Hydrogen mass repartitioning [63] was employed, enabling integration timestep of 8 and 4 fs for long- and short-range interactions respectively using the r-RESPA multiple time-stepping algorithm [64]. Each tetramer system was energy-minimized for 1,000 steps, followed by 5 ns of equilibration with soft harmonic restraints applied to all heavy protein atoms. A subsequent 10 ns equilibration phase was performed with restraints applied only to backbone heavy atoms. Production simulations were performed under two different conditions for 1 µs: (i) with all restraints removed, or (ii) with the four terminal Ala90 residues restrained at their initial positions using soft harmonic restraints applied to their backbone atoms (serving as a proxy for the full-length proteins anchored in the membrane). In (i), the overall rotational motion of the complexes was prevented (to account for the anisotropic shape of the rectangular simulation box) by applying a harmonic restraint to an orientational collective variable defined in the COLVAR module [65]. Specifically, the orientation component of the COLVAR library was employed, using all backbone atoms of the proteins as the reference atom group. This restraint preserved the global orientation of the complex while allowing internal conformational flexibility.

All analyses were performed using VMD [66]. Root-mean-square deviations (RMSD) were calculated relative to the initial structures using backbone atoms, considering either the entire protein or specifically the tetramerization interface (residues 120–156). The angle formed by the two dimers arranged in a head-to-head configuration (see Fig. 5, panel d) was quantified as the angle between two reference vectors, each associated with one dimer. For each dimer, the reference vector was defined as connecting the center of mass of the two Cα atoms of residue 100 to the center of mass of the two Cα atoms of residue 125.

Molecular dynamics simulations of the truncated ATP synthase *b* tetramer (residues 89–156) were used to reconstruct a dynamic model of the corresponding full-length tetrameric assembly (see Fig. 6, Panel a). For each trajectory frame, the structure of the complete *b* dimer observed in the *Escherichia coli* ATP synthase structure (PDB ID: 5T40) was systematically superimposed onto each truncated dimer of the simulated tetramer. The superposition was performed using backbone atoms of the overlapping region (residues 120-140). This procedure generated a frame-resolved structural model of the full-length tetramer, including the transmembrane segments, arranged in a head-to-head configuration.

The membrane-emergent residue was identified from independent simulations of the full-length dimer embedded in a lipid bilayer (see Supplementary Fig. 4). For each residue within the interfacial region (residues 30 to 44), the maximal projection along the membrane normal (z-axis) sampled during the trajectory was calculated and compared to the maximal z positions of the lipid polar headgroups. The residue ASP42, whose maximal z coordinate most closely matched the lipid headgroup boundary, was defined as the membrane–water interface residue.

Using the reconstructed dynamic model of the full-length tetramer, the distribution of membrane-to-membrane distances in the head-to-head assembly was estimated. This distance was defined as the separation between the centers of mass of the Cα atoms of residue Asp24 from each of the two opposing dimers, providing a geometric proxy for the intermembrane spacing imposed by the tetrameric arrangement (see Fig. 6, Panel c and Supplementary Fig. 4).

## Acknowledgements

This work was supported by CNRS, Université Paris Cité, LabEx DYNAMO (ANR-LABX-011), EQUIPEX (CACSICE ANR-11-EQPX-0008), and CPER-équipement PSL-RESOLUTION, notably through funding of the Proteomic Platform of IBPC (PPI). The authors acknowledge support from the Agence Nationale de la Recherche (ANR) through the funding of GenCaps (ANR-17-CE09-0007-02) and FLIPOSOME (ANR-20-CE06-0007) projects to C.T. and B.M.. M.K. is jointly supported by LabEx DYNAMO and the project COFUND FP-DYNAMO PARIS, which has received funding from the European Union’s Horizon 2020 research and innovation programme under the Marie Sklodowska-Curie grant agreement No 101034407. F.W. thanks Prof. Dr. Benjamin D. Engel for hosting him at the Biozentrum, supported by the Swiss National Science Foundation (SNSF) with an Ambizione Grant (number 216094). We thank Kathrin Marheineke for the critical reading of the manuscript, Marc Uzan for fruitful discussions and Oussama Houha for technical help. F.A. is supported by a French Ministry of Higher Education, Research, and Innovation PhD fellowship. J.R is supported by GenCaps and LabEx DYNAMO. We thank Biomics Platform, C2RT, Institut Pasteur, Paris, for RNA sequencing, which is supported by France Génomique (ANR-10-INBS-09) and IBISA.

## Declarations

- Competing interests. The authors declare no competing interests.
- Data availability. Cryo-EM tomograms are available at EMDB (accession code: EMD-56902, EMD-56903, EMD-56904, EMD-56905). Mass spectrometry data are available at MassIVE (accession code: PXD074300). RNA sequencing data are deposited at GEO, with accession number GSE320264. AlphaFold2 models and MD trajectories are available from the corresponding authors upon reasonable request. Source data for all graphs are provided with this paper.
- Expression plasmids and host are available from the corresponding authors upon request.
- Code availability
- Author contributions. B.M., F.Z., F.W., M.K., J.R., C.T., and F.D. conceived and designed the experiments. M.K., J.R., F.A., F.S., F.B., M.H., P.T., O.I., C.M., B.M., and F.Z. performed the experiments and contributed to data analysis. B.M. and F.Z. wrote the manuscript with contributions from all authors.

