## supplementary data for "F_1_F_0_-ATP-synthase subunit *b* head-to-head interactions shape intracellular membranes in *Escherichia coli*"

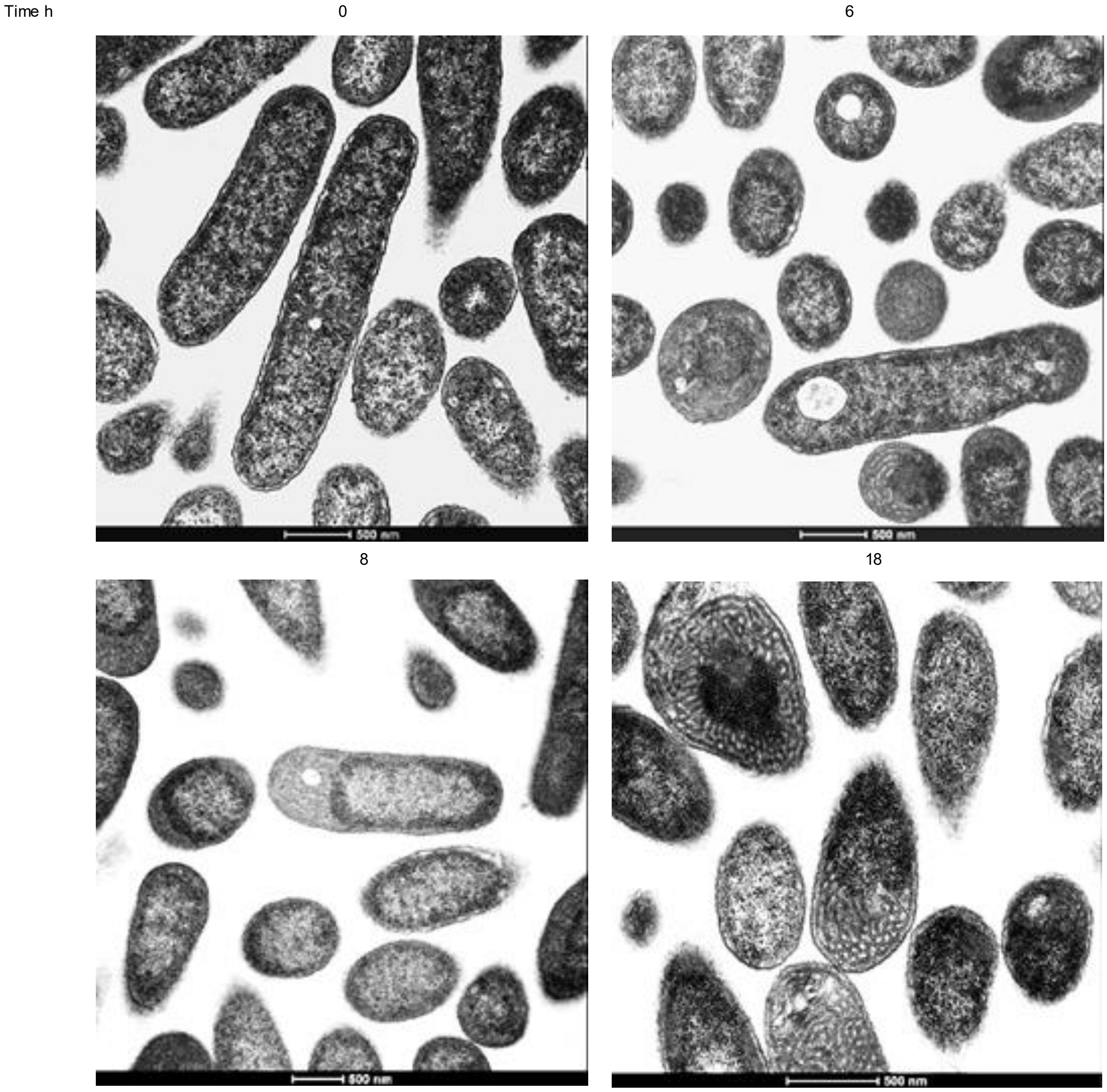


**Supplementary Fig 1: Kinetics of membrane proliferation**.

Transmission electron microscopy images of *E. coli* C43(DE3) cells overproducing ATP synthase subunit *b* taken over a time course after induction.

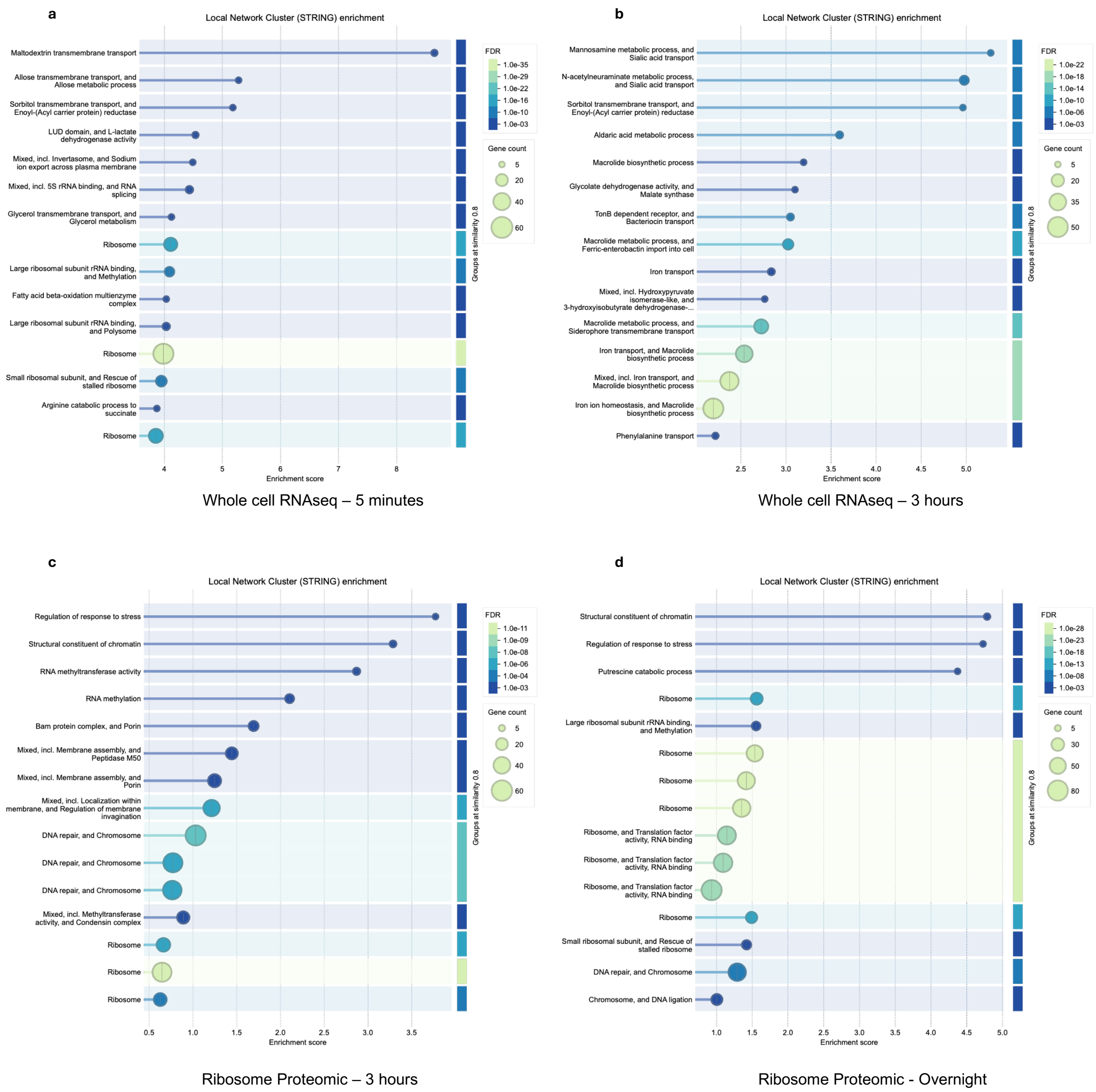


**Supplementary Fig 2: Comparing functional profiles among RNAseq and ribosomal proteomics data using STRING software.**

The four experimental clusters being compared are:

**a**, RNAseq 5 minutes.

**b**, RNAseq 3 hours.

**c**, ribosomal proteomics 3 hours.

**d**, ribosomal proteomics overnight.

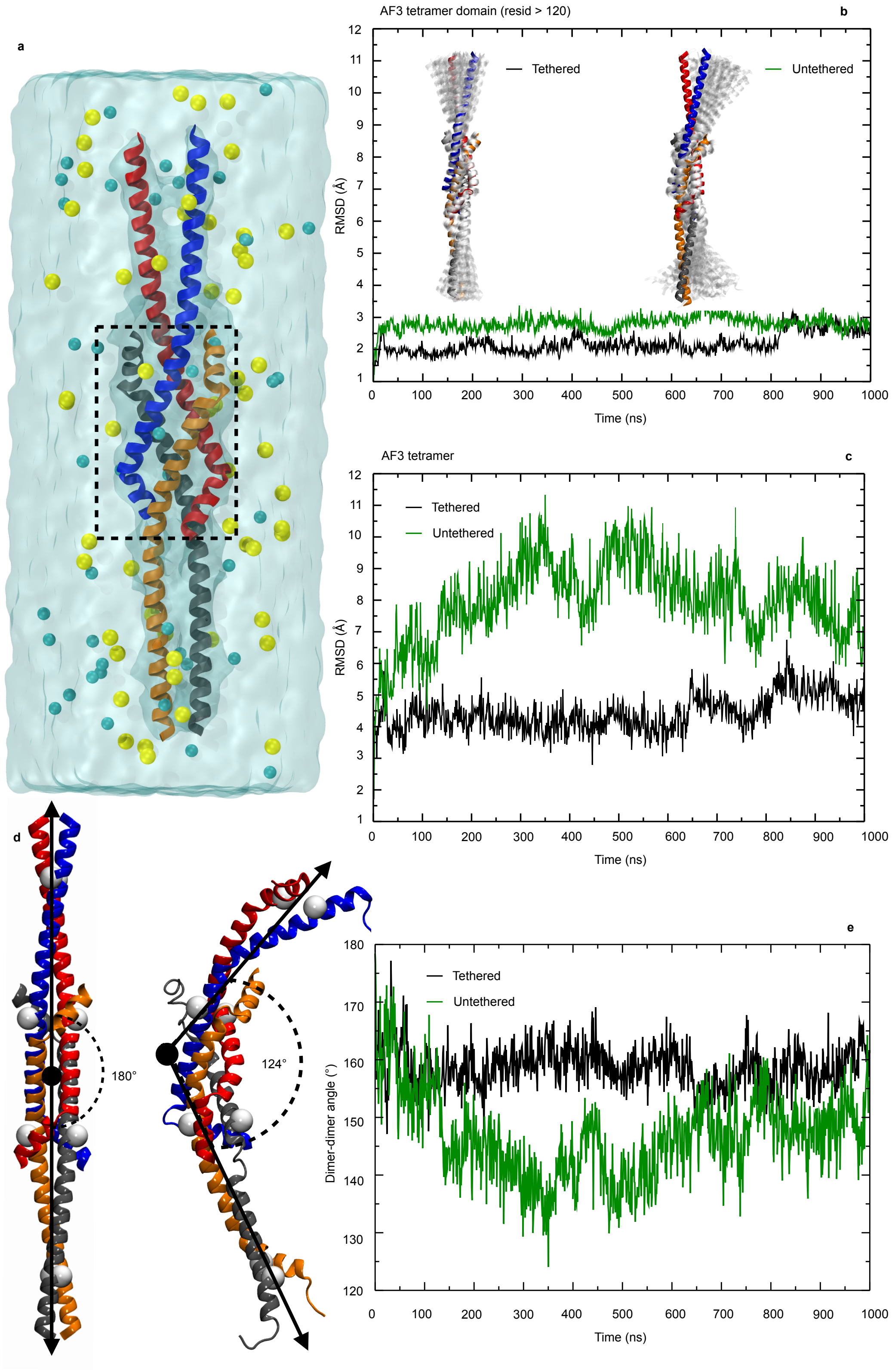


**Supplementary Fig. 3: Structural dynamics of an AlphaFold3-predicted tetramer of the *E. coli* ATP synthase subunit *b* (residues 90–156). (a)** Molecular dynamics (MD) simulation setup of the AF3 tetramer solvated in an explicit water box (light blue transparent surface)

containing 0.150 M NaCl ($Na^+$ and $Cl^-$ ions shown as yellow and cyan spheres, respectively). The dashed rectangle highlights the tetramerization domain (residues 120–156). Time evolution over 1 µs MD trajectories of the backbone root-mean-square deviations (RMSD) for **(b)** the tetramerization domain and **(c)** the full AF3 tetramer. Black curves correspond to a trajectory in which the tetramer is tethered (the four terminal Ala90 residues are harmonically restrained to their initial positions), whereas green curves correspond to a trajectory in which the tetramer is untethered. Insets in panel **(b)** show 50 conformations (transparent white cartoons) sampled along the MD trajectories for the tethered (left) and untethered (right) tetramers, overlaid on the initial AF3 model (solid-colored cartoons). **(d)** Tetramer structures representing the extreme angles between the two-forming dimers, sampled along MD trajectories. Arrows indicate the axes used to measure the angle, and white spheres mark the Cα atoms defining these axes. **(e)** Time evolution of the angles formed by the two dimers for tethered (black line) and untethered (green line) tetramers.

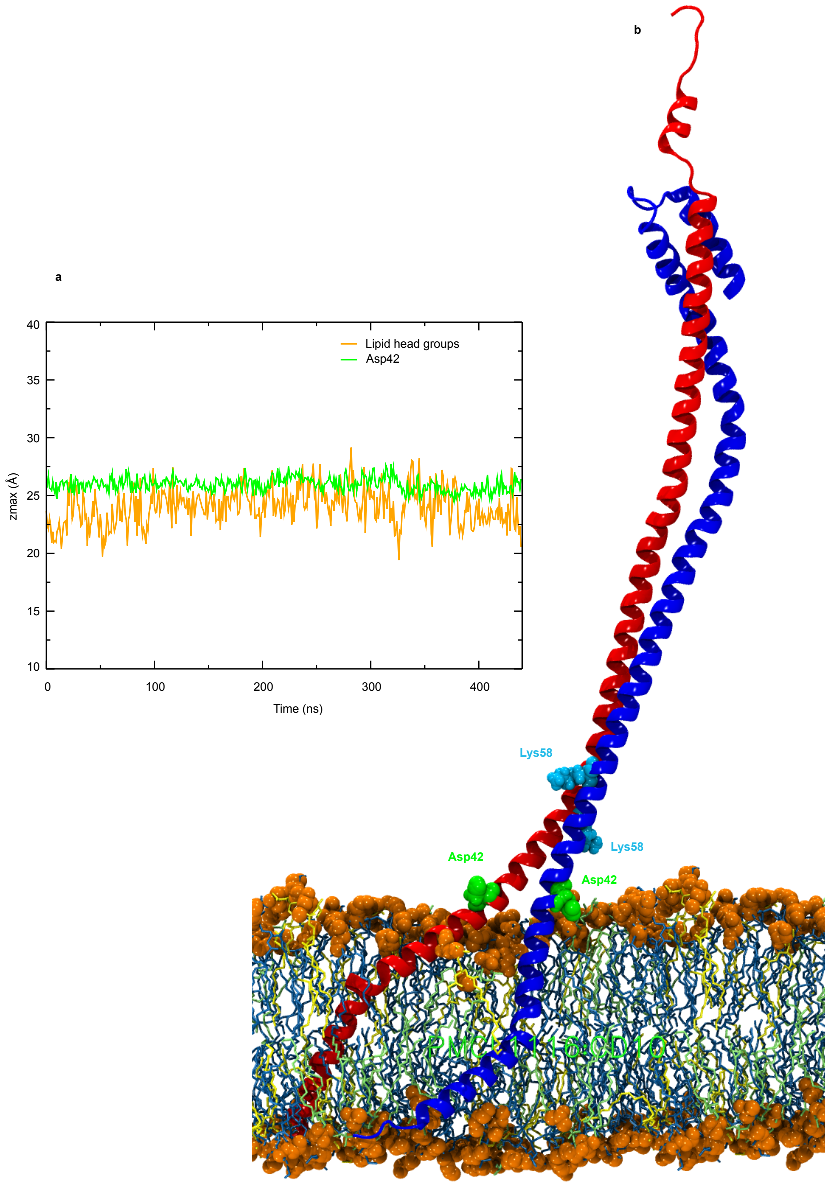


**Supplementary Fig. 4: Dynamics of the full-length *E. coli* ATP synthase subunit *b* dimer (PDB:5T4O). (a)** Time evolution of extremal positions of the atoms of lipid headgroups (orange line) and Asp42 (green line) along the axis orthogonal to the bilayer plane. **(b)** Snapshot taken along a 450 ns MD trajectory of the b subunit dimer embedded in a heterogeneous bilayer composed of POPG, POPE and cardiolipins (yellow, blue and green licorices respectively). Orange spheres represent the lipid polar headgroups (ions and water are omitted for clarity). Asp42 residues are represented by van der Waals green spheres. Lys58 (first trypsin cleavage site) is shown as van der Waals blue speres.
